# *Drosophila* female-specific brain neuron elicits persistent position- and direction-selective male-like social behaviors

**DOI:** 10.1101/594960

**Authors:** Yang Wu, Salil S. Bidaye, David Mahringer

## Abstract

Latent neural circuitry in the female brain encoding male-like mating behaviors has been revealed in both mice and flies. In *Drosophila*, a key component of this circuitry consists of the *doublesex*-expressing pC1 neurons, which were deemed to exist in both sexes and function based on the amount of cells being activated. Here, we identify pC1-alpha, a female-specific subtype of pC1, as responsible for inducing persistent male-like social behaviors in females. We demonstrate that activation of a single pC1-alpha neuron is sufficient for such induction in a position- and direction-selective manner, and activity of pC1-alpha neurons as a whole is indispensable for maintaining normal sexual receptivity. These dual functions of pC1-alpha may require different neurotransmission, with acetylcholine specifically required for the former but not the latter. Our findings suggest that pC1-alpha may be the female counterpart of male P1 due to their shared similarities in morphology, lineage, and social promoting function.

## Introduction

The neural substrates of social behaviors, such as courtship and aggression, and their dimorphisms between the sexes have been studied extensively in *Drosophila melanogaster* (Anderson, 2016; Auer and Benton, 2016; Dickson, 2008). Like most unisexual species shaped by sexual selection, flies exhibit conventional sex roles: males compete with each other via aggression, but perform an elaborate ritual of courtship toward choosy females, who rarely fight and almost never court (Bateman, 1948; Trivers, 1972). Such sexually distinct social behaviors are neurally hardwired through the coordinated actions of the sex determination genes *doublesex* (*dsx*) and *fruitless* (*fru*) (Demir and Dickson, 2005; Kimura et al., 2005; Manoli et al., 2005; Rideout et al., 2010; Stockinger et al., 2005). *Dsx*/*fru*^+^ P1 neurons promote an internal state that enhances both courtship and aggression (Bath et al., 2014; Hoopfer et al., 2015; Inagaki et al., 2014). Their presence in males and absence in females due to developmental apoptosis may explain the sexually dimorphic patterns of flies’ social behaviors (Kimura et al., 2008). Yet, recent studies in both mice and flies have revealed the existence of latent neural circuitries that encode male-like mating behaviors in the female brain (Kimchi et al., 2007; Rezaval et al., 2016; Wei et al., 2018), suggesting a conserved bisexual nature of brain organization and function (Dulac and Kimchi, 2007). In flies, this circuitry consists of the *dsx*^+^ pC1 cluster of neurons that are also essential for female sexual receptivity (Rezaval et al., 2016; Zhou et al., 2014). However, the pC1 neurons and its subsets were generally treated as more or less homogeneous and present in both sexes (Rezaval et al., 2016; Zhou et al., 2014), and their function in triggering male-like courtship behavior was deemed to depend on the amount of cells being activated (Rezaval et al., 2016). Here, we identify pC1-alpha, a female-specific subtype of the pC1 cluster, as responsible for inducing persistent male-like social behaviors in females. We demonstrate that activation of a single pC1-alpha neuron is sufficient for such induction in a position- and direction-selective manner, and activity of pC1-alpha neurons as a whole is indispensable for maintaining normal female sexual receptivity. These dual functions of pC1-alpha may require different neurotransmission, with acetylcholine specifically required for the former but not the latter. Our findings suggest that pC1-alpha may be the female counterpart of male P1 due to their shared similarities in morphology, lineage, and social promoting function, and may likely represent a novel cell type that arises specifically in the female brain from cooption of P1-like neural templates.

## Results

### Thermogenetic screen and MARCM identified two cell types for inducing male-like social behaviors

We performed an unbiased large-scale thermogenetic screen in flies (Bidaye et al., 2014), which utilized the Vienna Tiles (VT) collection of *GAL4* driver lines (Tirian and Dickson, 2017) and a *UAS-trpA1* transgene to target expression of the thermosensitive cation channel TrpA1 (Hamada et al., 2008) to random but defined subsets of neurons. *VT-GAL4 UAS-trpA1* flies were slowly warmed to ~32°C to open the TrpA1 channel and visually examined for ectopic behaviors. We screened 3470 lines and identified six, in which specifically females showed male-like social behaviors toward other flies at elevated temperatures but not at the control temperature of ~25°C (Movie S1). These male-like behaviors included courtship and aggression by a TrpA1-expressing test fly toward a non-confronting and a confronting control or test fly, respectively (Movie S1). When multiple TrpA1-expressing test females were co-housed in the behavior arena, in addition to courtship and aggression, we also observed chaining behavior (Movie S1). The courtship-like behavior mainly consists of frequent orientation and following and occasional tapping and unilateral wing extension (Movie S2). Such courtship-like behaviors often elicited ovipositor extrusion from female targets (Figure S1A), but we recorded no courtship song (Movie S2; Figure S1B and C). The most robust of these induced male-like social behaviors is the following behavior (including orientation), which is crucial for initiation and maintaining proximity for subsequent social actions. Therefore, we focused our analysis on the induced following behavior in the females of these lines.

The expression patterns of the six identified VT lines, examined with a membrane-tethered mCD8-GFP (green fluorescent protein) reporter, showed that a female-specific cell type might be present in all of them (Figure 1A). This cell type, consisting of only two to three neurons in each hemisphere, morphologically resembles *dsx*^+^ pC1 (Rezaval et al., 2016; Zhou et al., 2014). To achieve more restricted expression, we applied MARCM (mosaic analysis with a repressible cell marker) (Lee and Luo, 1999) on the sparsest *VT2064-GAL4* line to generate mosaic females, in which expression of mCD8-GFP and TrpA1 were targeted to random subsets of neurons labeled in the line. We then monitored them for following behavior in warmed chambers and determined their exact expression patterns. We observed induced following behavior in 37 flies, expression analysis of which, in comparison to 34 flies that did not show the phenotype, revealed that induction of following behavior strongly correlates with expression in two cell types (Figure 1B to D; Figure S1G). Cell type 1, located in the dorsal posterior brain, is the shared female-specific pC1-like neuron, whose neurites project first anterodorsally to arborize extensively toward the lateral and medial protocerebrum regions and then transversally to contralateral regions (Figure 1E). Cell type 2 is a nearby local interneuron innervating the central complex (Figure 1F). Their enrichment in females showing induced following behavior was unlikely due to biased MARCM labeling, as comparable numbers of cell types were labeled in non-following females (Figure S1D). We conclude that activation of cell type 1 and/or 2 elicits male-like social behaviors in females. However, we hypothesize that only cell type 1 is responsible, because there are no MARCM flies that only label cell type 2 but show the behavior and not all six positive VT lines label cell type 2 (Figure 1A, C and D; Figure S1E). Cell type 2 may be linked to cell type 1 clonally, as they were always colabeled ipsilaterally (Figure S1F).

**Figure 1.**
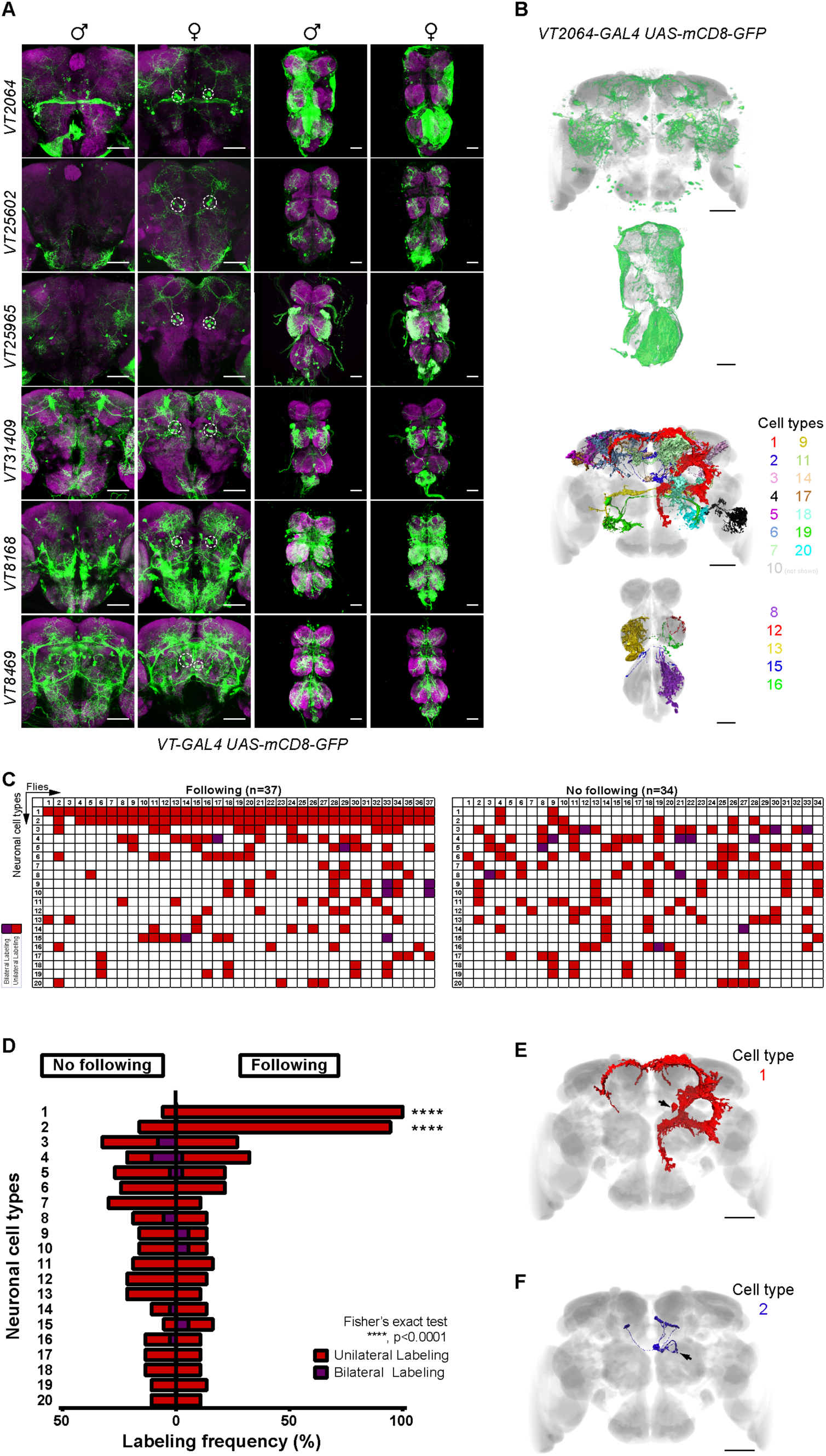
Thermogenetic screen and MARCM identified two cell types for inducting male-like social behaviors. (A) Brains (left) and ventral nerve cords (VNCs, right) of the six identified GAL4 lines expressing mCD8-GFP, stained with GFP antibody (anti-GFP) (green) and nc82 (magenta). Dashed circles indicate a shared female-specific neuron. (B) Overall expression (top) and individual segmentations of 20 most frequently labeled cell types (neuron 10 not shown, bottom) in *VT2064.GAL4* MARCM females, following non-rigid registration onto common reference templates. A total of 57 cell types were observed (37 in the brain and 20 in the VNC). (C) Expression patterns of MARCM females, showing following behavior (n=37, left) or no following (n=34, right) upon activation. Each row represents scores for a cell type, and each column a single fly. Unilateral and bilateral labeling were indicated by red and purple colors, respectively. (D) Labeling frequency for each cell type in MARCM females with and without induced following behavior. ****p<0.0001, Fisher’s exact test. (E and F) Registered single-neuron segmentations of cell type 1 (E) and 2 (F). Arrows indicate cell body locations. Scale bars, 50 µm. See also Figure S1 and Movie S1 and S2.

### Sparse genetic reagents confirmed that activation of cell type 1 elicited male-like social behaviors

From the six positive GAL4 lines, we generated corresponding split-GAL4 hemi-drivers to be combined pair-wise (Luan et al., 2006; Pfeiffer et al., 2010) and Hap1 lines using an alternative yeast-derived Hap1-HBS expression system (M.H. & B.J.D., personal communication). We derived a much sparser set of four split-GAL4 combinations (referred to as *Split-1 to 4-GAL4*, see method for exact genotypes) and one Hap1 driver (*VT8469-Hap1*), all of which labeled only cell type 1 but not cell type 2 (Figure 2A and B; Figure S2A to C). Cell type 1, with 2 cells in each hemisphere, is the only overlap between *Split-1-GAL4* (*VT25602.p65ADZp; VT2064.ZpGAL4DBD*) and *VT8469.Hap1*, each of which labels only one additional cell type in the brain (Figure 2A, B and H). Both of these drivers triggered no obvious behaviors in isolated females and male-like social behaviors in females individually paired with a target fly at both 28°C and 31°C (Figure 2C and D; Movie S3), independent of target fly sex (Figure 2E) and their own mating status (Figure 2F). Remarkably, induced following behavior persisted, even 15 min after returning to 25°C from 10-min activation at 31°C (Figure 2G). Using *VT8469.Hap1*, we found that cell type 1 is colabeled by *dsx-GAL4* (Figure 2I), together with its cell body position, confirming that it indeed belongs to the *dsx*^+^ pC1 cluster. Its morphology, revealed by these sparse drivers and single MARCM clones, overall resembles that of pC1 cluster, as previously defined by *71G01-LexA*∩*dsx-GAL4*, but clearly differs from those revealed by multicolor flip-out analysis (Zhou et al., 2014). Its cell number (1.91±0.20, n=16) is fewer than what has been reported for pC1 (8.0±2.0) or its *71G01-LexA*∩*dsx-GAL4* defined subset (6.6±0.3) (Zhou et al., 2014). We also confirmed that it was labeled by all the screen-identified GAL4 lines, by examination of derived split-GAL4 combinations (Figure S2A to C) or the original positive GAL4 lines colabeled with *VT8469.Hap1* (Figure S2D). These data suggest that cell type 1 is likely a female-specific stereotypic subtype of *dsx*^+^ pC1. Therefore, we refer to it as pC1-alpha for its ability to induce male-like behaviors and conclude that the female-specific pC1-alpha is functionally responsible for inducing persistent male-like social behaviors in females.

**Figure 2.**
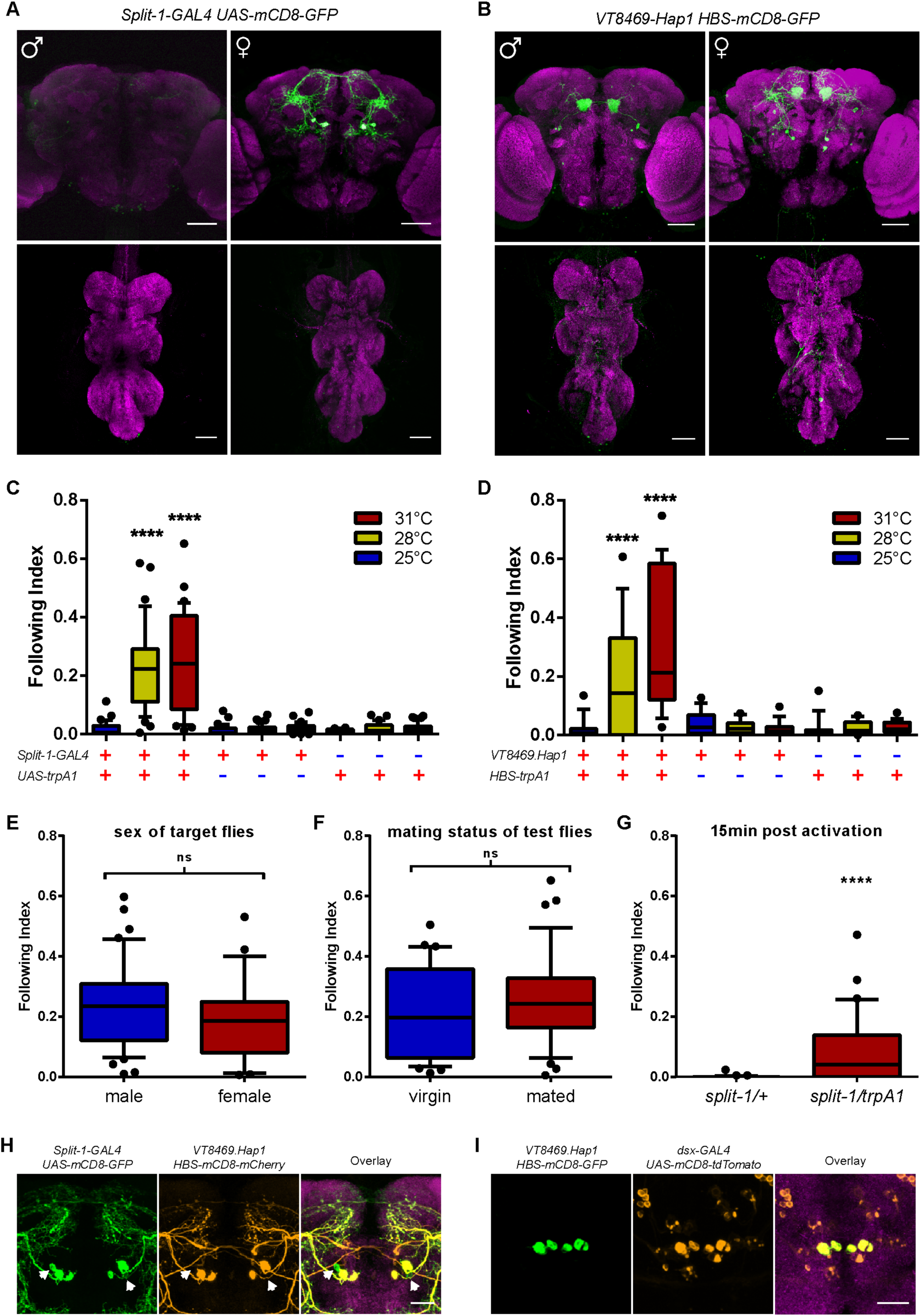
Sparse genetic reagents confirmed that activation of cell type 1 elicited male-like social behaviors. (A and B) Posterior view of brain (left) and VNC (right) of males and females carrying *Split-1-GAL4 UAS-mCD8-GFP* (A) and *VT8469.Hap1 HBS-mCD8-GFP* (B), stained with anti-GFP (green) and nc82 (magenta). (C and D) Percentage of time females with indicated genotypes spent in following during 10-min assays individually paired with a wildtype fly under different temperatures (n=25-45 (C) or 15-26 (D)). (E to G) *Spilt-1-GAL4 UAS-trpA1* females induced male-like social behaviors, which were independent of sex of target flies (E, n=41 and 26) and their own mating status (F, n=36), and persisted, even 15 min after returning to 25°C from 10-min incubation at 31°C (n=31-33). (H and I) Brains of females carrying *Split-1-GAL4*, *UAS-mCD8-GFP*, *VT8469*.*Hap1*, *HBS*-*mCD8*-*mCherry* (H) and *VT8469*.*Hap1*, *HBS-mCD8-GFP*, *dsx-GAL4*, *UAS-mCD8-tdTomato* (I), stained with anti-DsRed (oragne), anti-GFP (green), and nc82 (magenta). A total of four pC1-alpha neurons per brain were co-labeled among the two drivers and *dsx-GAL4*. Another two neurons in the vicinity were only weakly labeled by *Split-1-GAL4* (H, arrows). Whiskers represent 10-90 percentiles; ****, p<0.0001, Mann-Whitney test. Scale bars, 50 µm (A and B) and 20 µm (H and I). See also Figure S2 and Movie S3.

### Single pC1-alpha neurons induce position- and direction-selective male-like following with directional neural signals

During the *VT2064.GAL4* MARCM experiment, we noticed that one of the mosaic females specifically labeled a single pC1-alpha neuron in the brain (Figure 3A) and directed its induced male-like following behavior only toward target flies moving in a specific direction (Fly 1; Figure 3A; Movie S4). To confirm this observation, we performed a second MARCM experiment using flies carrying *VT8469.Hap1 HBS-GAL4* transgenes, effectively converted into a GAL4 driver. We obtained another four flies that labeled one or two ipsilateral pC1-alpha neurons (Fly 2 to 5; Figure 3B). In these flies, male-like following was often discontinuous, fragmented, and only observed when target flies moved across their field of view in contralateral-to-ipsilateral directions: Fly 1 with left pC1-alpha activated followed mainly toward flies moving in front of her from right to left, while Fly 2 to 5, which labeled the right pC1-alpha, showed following behavior toward flies moving from left to right (Figure 3C, D and F; Movie S4). This was in direct contrast to flies with bilateral pC1-alpha neurons activated, who showed longer bout duration and pursued target flies not only moving across their field of view from both directions, but also those directly in front of them (Figure 3D and G; Movie S4). We used MateBook (Ribeiro et al., 2018) to extract positional and angular information of both test and target flies (Figure 3E to G) and manually annotated video recordings for frames where test flies showed following behavior (Movie S4). We compiled frames of about 20 following bouts from each of the five MARCM flies (with direction for Fly 1 reversed), together with those from five *Split-1 UAS-trpA1* flies, and examined the distribution of distance and relative angles Θ_01_ and Θ_Heading_ between test and target flies (Figure 3E). Distance was similarly biased toward small values for both MARCM and *Split-1* flies, but angles Θ_01_ and Θ_Heading_ were specifically biased to the ipsilateral side for MARCM flies (Figure 3H) and to both sides for *Split-1 UAS-trpA1* flies (Figure 3I). We conclude from these data that activation of a single female pC1-alpha neuron is sufficient to induce male-like following behavior in a position- and direction-selective manner with a bias toward the ipsilateral side.

**Figure 3.**
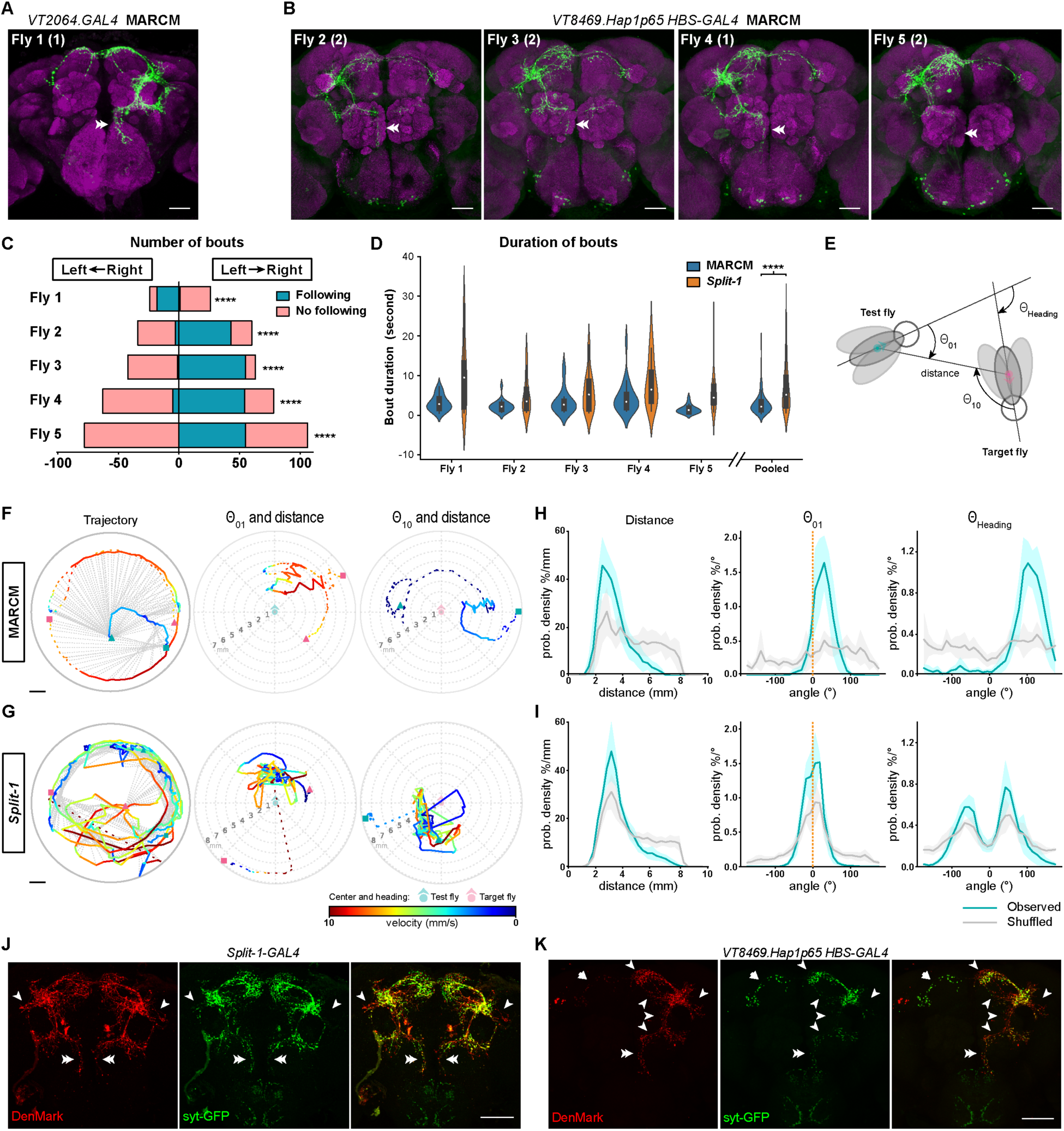
Single pC1-alpha neurons induce position- and direction-selective male-like following with directional neural signals. (A and B) Brains of MARCM females using *VT2064.GAL4* (A) or *VT8469.Hap1 HBS-GAL4* (B) that labels one or two ipsilateral pC1-alpha neurons (as indicated in the parenthesis), stained with anti-GFP (green) and nc82 (magenta). (C) Bout frequency showing following (blue) or no following (pink) behavior, when single-paired target flies moved across the visual fields of test flies from right-to-left or left-to-right direction. ****p<0.0001, Fisher’s exact test. (D) Duration of following bouts in MARCM and *Split-1-GAL4* flies expressing TrpA1 in pC1-alpha, when individually paired with a target fly (n=19-21 bouts for each fly). ****p<0.0001, Mann-Whitney test. (E) Distance and angle θ_01_, θ_10_, and θ_Heading_ between test female and target fly. (F and G) Tracked trajectory (left) and polar plots, showing angle θ_01_ and distance to target fly from the perspective of test female fly (middle) and angle θ_10_ and distance to test female from the perspective of target fly (right), of a following bout of MARCM Fly 2 (F) and a *Split-1-GAL4* fly (G) expressing TrpA1 in pC1-alpha. For trajectory plots, scale bars are 1mm, circular outlines represent 10mm arenas, and dotted lines connect corresponding fly positions within the same frame. For polar plots, 0° is at the top and degrees increase clockwise with a range of (−180°,180°]. Color codes represent speed of corresponding fly in mm/s. (H and I) Probability density functions of distance, angle θ_01_ and θ_Heading_ between test females of indicated genotype and single-paired target flies, for observed (blue) and shuffled (gray) data. Shaded areas represent standard deviation. (H) MARCM Fly 1-5, with direction for Fly 1 reversed. (I) *Split-1-GAL4 UAS-trpA1*, n=5. (J and K) Brains of females expressing polarity markers, DenMark and syt-GFP, in all (J) or one (K) pC1-alpha neurons, stained with anti-DsRed (red) and anti-GFP (green). Arrows and arrowheads indicate predominant dendritic and pre-synaptic regions, respectively. Scale bars, 50µm (A, B, J, and K) and 1mm (F and G). Double arrowheads indicate the medial ventral neurites of pC1-alpha. See also Movie S4.

Such ipsilateral bias led us to examine the information flow in the complex arborizations of pC1-alpha neurons. Expression of a somato-dendritic marker DenMark (Nicolai et al., 2010) and a presynaptic marker synaptotagmin-GFP (Zhang et al., 2002) in all pC1-alpha neurons showed an almost complete overlap of the two markers (Figure 3J). However, when expressed in only one or two ipsilateral cells in mosaic females generated by MARCM, the two markers partially separated. Namely an exclusively dendritic ipsilateral ventral and an exclusively synaptic contralateral dorsal region, while the ipsilateral dorsal region displayed both markers (Figure 3K, n=8). Such a pattern may suggest an ipsilateral-to-contralateral and ventral-to-dorsal flow of information, which may be modulated in between.

### pC1-alpha requires acetylcholine for following behavior induction but not normal sexual receptivity

We subsequently examined which neurotransmitter pC1-alpha neurons used for induction of male-like social behaviors. Antibody staining against choline acetyltransferase (*ChAT*), which catalyzes the biosynthesis of the neurotransmitter acetylcholine (ACh), showed co-localized signals with pC1-alpha neurons, suggesting that they are cholinergic (Figure 4A). In addition, when we knocked down *ChAT* specifically in pC1-alpha neurons, by expressing two different RNAi lines with *Split-1-GAL4*, induction of male-like behavior was significantly reduced or abolished (Figure 4B). We conclude that pC1-alpha neurons induce male-like following behavior via ACh.

**Figure 4.**
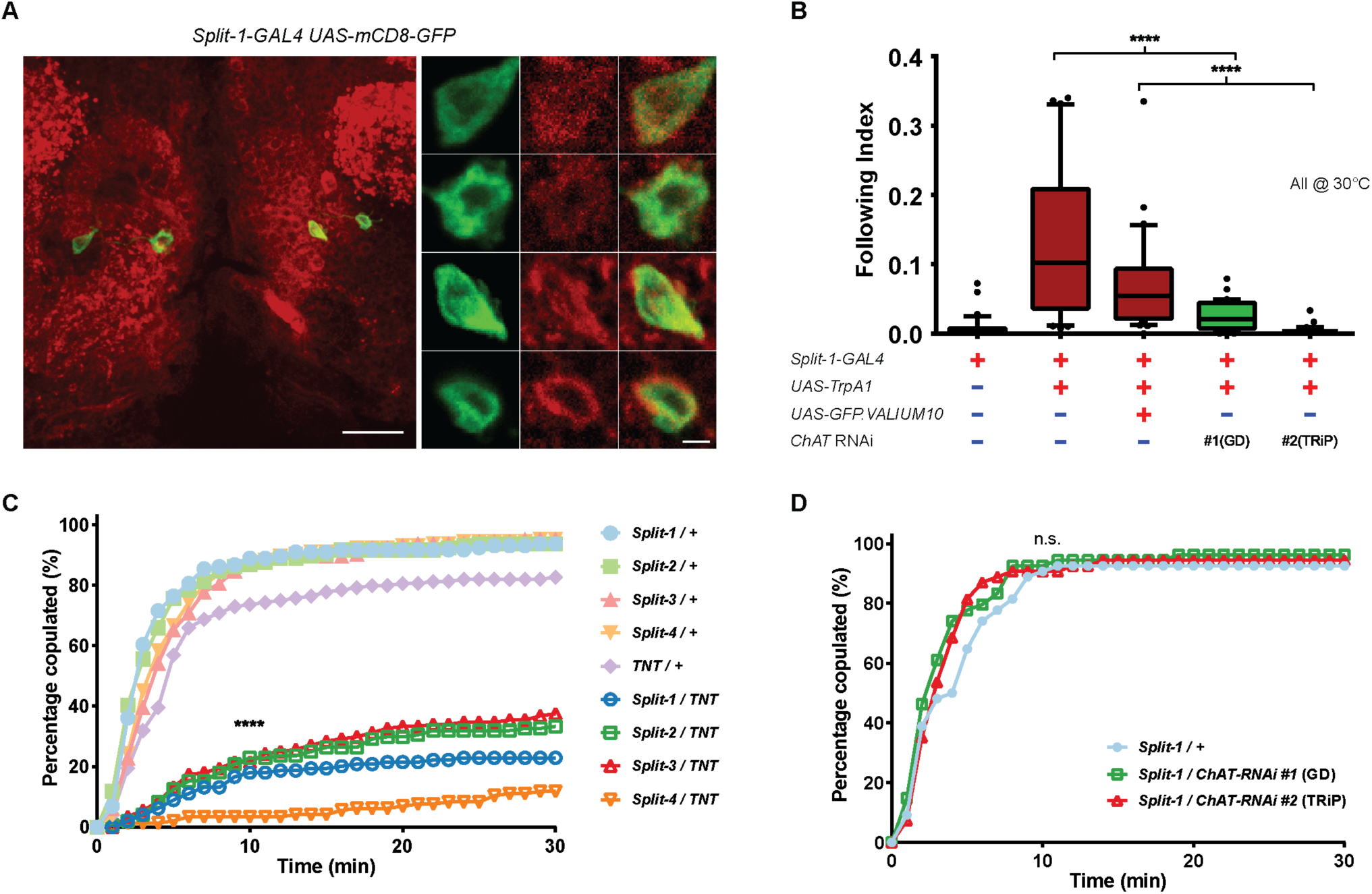
pC1-alpha requires acetylcholine for following behavior induction but not normal sexual receptivity. (A) Brain of a *Split-1-GAL4 UAS-mCD8-GFP* female, at low (left) and high (right) magnification, stained with anti-ChAT (red) and anti-GFP (green). Scale bars, 20µm (left) and 2µm (right). (B) RNAi knock-down of *ChAT* in pC1-alpha blocked induction of male-like following behavior at ~30°C. Percentage of time flies with indicated genotypes spent in following target flies during 10-min assays (n=29-33). *UAS-GFP.VALIUM10* was used as control for TRiP *ChAT* RNAi line (#2). ****p<0.0001, Mann-Whitney test, whiskers represent 10-90 percentiles. (C) Inactivation of pC1-alpha neurons reduced sexual receptivity in females carrying *Split-1* to *4-GAL4 UAS-TNTE*. Data are represented as mean ± SD. ****p<0.0001 when compared to corresponding controls at 10min, Chi-square test. (D) RNAi knock-down of *ChAT* had no effect on female receptivity. Data are represented as mean ± SD. n.s., p>0.05 when compared to control at 10min, Chi-square test.

Since *dsx*^+^ pC1 neurons play an important role in female sexual receptivity (Rezaval et al., 2016; Zhou et al., 2014), we asked whether activity of the specific pC1-alpha subtype is essential for normal female receptivity, and if so, whether ACh is required for its function. We first inactivated pC1-alpha neurons by combining *Split-1* to *4-GAL4* with a *UAS-TNT* transgene, which encodes an inhibitor of synaptic transmission (Sweeney et al., 1995). We observed a dramatic reduction of copulation in comparison with the controls, suggesting an essential role of pC1-alpha in maintaining normal female receptivity (Figure 4C). However, when we expressed the two *ChAT* RNAi lines in pC1-alpha neurons, female receptivity remained comparable to controls (Figure 4D), suggesting a potential co-transmission mechanism (Kupfermann, 1991; Nusbaum et al., 2001). However, it is also possible that our RNAi lines may have reduced *ChAT* expression in pC1-alpha neurons to a level that could block the induction of male-like behavior, which requires stronger activation, but leave intact female receptivity functions, which may require weaker activation.

## Discussion

The *dsx*^+^ pC1 neurons, previously reported to constitute the latent courtship circuit in the brain of *Drosophila* females, were characterized as a functional unit that is sex-common and cell-number dependent in eliciting male-like courtship (Rezaval et al., 2016). Here, we identify the exact cell type that is responsible for triggering such male-like social behavior in females and show its female specificity and sufficiency in recapitulating the overall phenotypes observed with the pC1 cluster. Our results reveal that pC1 neurons, although all expressing *dsx*, are likely rather heterogeneous, and it is the cell identity rather than the cell number that determines their function. In our case, activation of a single pC1-alpha neuron and inactivation of the whole pC1-alpha cluster are sufficient to induce male-like social behaviors and suppress sexual receptivity, respectively.

Our findings suggest that activation of pC1-alpha neurons triggers a visually guided following behavior that encourages overall social interactions, rather than just male-like courtship behavior alone. The outcome of such increased social interactions may be either courtship or aggression, depending on social feedback (Movie S1 and S3). When a female is paired with a target fly, visual sensory inputs of the target are constantly processed, but normally do not trigger male-like following behavior in the female. Its induction by pC1-alpha activation suggests a link between motion detection and locomotion circuitries, which is likely gated by pC1-alpha activity. Motion of small objects, such as a fly, is detected by the LC10 visual projection neurons to direct courtship behavior in males and likely male-like social behaviors in females as well (Ribeiro et al., 2018). Given that LC10 neurons are not selective to movement direction (Ribeiro et al., 2018), the position- and direction-selectivity of unilateral pC1-alpha activation likely emerges downstream of LC10, possibly by potentiation of downstream descending neurons that lead to turning and forward walking. It would be interesting to identify the exact location where visual information is integrated with pC1-alpha gating signals.

The activity of pC1-alpha likely promotes an internal state of arousal or motivation, due to its induction of persistent social behaviors in females paired with a target and the lack thereof in isolated females. This, in addition to its morphology, *dsx* expression and sex-specificity, is highly reminiscent of male P1 (Anderson, 2016). P1 likely also shares the same DM4 lineage with pC1-alpha, based on their homologous morphology (Figure S4A to D) (Ren et al., 2016). However, unlike P1 which fates to die and is faithfully rescued by blocking apoptosis in females (Kimura et al., 2008), pC1-alpha may never exist in males, as blocking apoptosis throughout DM4 lineage development failed to rescue male *dsx*^+^ neurons with the characteristic medial ventral neurites of pC1-alpha (Figure 3A, B, J and K, Figure S2E to G, double arrowheads) (Ren et al., 2016). Thus, we speculate that pC1-alpha likely represents a female-specific “*de novo*” P1-like cell type, which may be a result of nature’s tinkering (Jacob, 1977). We further speculate that such neuronal organization may allow sufficient flexibility to encode new mating strategies during evolution (Donegan and Ewing, 1980), including the extreme cases of sex role reversal, where females compete for male mates (Kokko et al., 2006).

In summary, our results demonstrate that activation of a single female-specific *dsx*^+^ pC1-alpha neuron is sufficient to induce male-like social behaviors in a position- and direction-selective manner, and activity of pC1-alpha neurons as a whole is indispensable for maintaining normal sexual receptivity. Cholinergic neurotransmission of pC1-alpha neurons is specifically required for male-like behavior induction, but not sexual receptivity regulation. Our findings suggest a link between male- and female-specific higher-order neurons that both promote a persistent internal state to enhance social behaviors, identifying pC1-alpha as P1’s female counterpart likely co-opted from P1-like templates to regulate female sexual receptivity. Therefore, our study provides additional circuit hypotheses that can be tested in further studies.

## Acknowledgements

We thank M. Haesemeyer for development and characterization of Hap1-HBS system and general comments on manuscript, K. Feng for generating *HBS-GAL4* fly and Y. Ding for help on courtship song analysis. All of the experimental work described here was performed in the lab of Barry J. Dickson at IMP in Vienna, Austria and Janelia Research Campus in Ashburn, USA, with funding from Boehringer Ingelheim & Howard Hughes Medical Institute, respectively.

## Author contributions

Y.W., S.B., and D.M. conducted the experiments; Y.W. designed the experiments and wrote the paper.

## Methods

### Fly stocks

The VT GAL4 collection and Hap1 lines were constructed according to (Pfeiffer et al., 2008), using the *attP2* landing site (Groth et al., 2004). Split-GAL4 lines were generated according to (Pfeiffer et al., 2010) and inserted into either *attP2* or *attP40* (Groth et al., 2004). Both VT collection and split-GAL4 lines have been described in greater details (Tirian and Dickson, 2017). MARCM stocks: 1) *hs-FLP*, *FRT^G13^*, *tub-GAL80*; 2) *FRT^G13^*, *UAS-mCD8-GFP*; 3) *neoFRT^19A^*; 4) *hsFLP*, *tub-GAL80*, *neoFRT^19A^* were obtained from Bloomington *Drosophila* Stock Center. Other stocks used were *UAS-trpA1* (von Philipsborn et al., 2011), *UAS-synaptotagmin-GFP* (Zhang et al., 2002), *UAS-DenMark* (Nicolai et al., 2010), *UAS-mCD8-tdTomato* (Toda et al., 2012), *ChAT*-RNAi-#1(GD) (Dietzl et al., 2007), and *ChAT*-RNAi-#2(TRiP) (Perkins et al., 2015). *HBS-GAL4*, *HBS-mCD8-mCherry*, *HBS-trpA1* were provided by M.H. and B.J.D.

### Behavioral assay and analysis

For activation with TrpA1, flies were raised at 22°C and collected shortly after eclosion, then aged in groups of 15-20 in separate vials for 10-14 days at 22°C before testing. During testing, flies were paired with *white-* wild type flies or TrpA1-expressing test flies and assayed for following behavior in 10mm chamber for 10min under different temperatures. Ambient temperatures were adjusted and maintained constant, using a heating glass ceiling and a power supply with a feedback controller. Percentage of time spent in following within a 10min assay, the following index, was used as a measure of following behavior. Videos were either automatically tracked using Matebook (Ribeiro et al., 2018) (Figure 2C to F, 4B) or manually analyzed (Figure 2G, 3C and D, 4C and D), or both (Figure 3F to I).

### MARCM (mosaic analysis with a repressible cell marker)

Flies carrying appropriated transgenes were crossed and progenies from the crosses were heat shocked at late embryonic and early larval stages for 1-1.5hr. Several MARCM experiments were conducted in this study: 1) For MARCM with *VT2064.GAL4*, flies carrying *FRT^G13^*, *UAS-mCD8-GFP*; *UAS-trpA1* and *hs-FLP*; *FRT^G13^*, *tub-GAL80*; *VT2064.GAL4* were crossed. 2) For MARCM with *VT8469.Hap1*, *neoFRT^19A^*; *HBS-GAL4*; *VT8469.Hap1* flies were crossed to *hsFLP*, *tub-GAL80*, *neoFRT^19A^*; *UAS-mCD8-GFP*; *UAS-TrpA1*. 3) For MARCM with *VT8469.Hap1* and two polarity markers, *neoFRT^19A^*; *HBS-GAL4*; *VT8469.Hap1* and *hsFLP*, *tub-GAL80*, *neoFRT^19A^*; *UAS-DenMark*, *UAS-synaptotagmin-GFP*; *UAS-DenMark*, *UAS-synaptotagmin-GFP* were used.

### Immunohistochemistry and imaging

Immunohistochemistry was performed as described in (Yu et al., 2010). Confocal stacks were acquired on a Zeiss LSM 700 and we used rabbit anti-GFP (1:6000, Torrey Pines Biolabs), chicken anti-GFP (1:10000, Abcam), rabbit anti-DsRed (1:1000, Clontech), mouse mAb anti-ChAT (1:100, DSHB) and secondary Alexa 488, 568, and 647 (1:1000, Invitrogen) antibodies. Maximum intensity projections were generated using ImageJ.

### Split-GAL4 genotypes

Split-1-GAL4: VT25602.p65ADZp@attP40; VT2064.ZpGAL4DBD@attP2

Split-2-GAL4: VT2064.p65ADZp@attP40; VT31409.ZpGAL4DBD@attP2

Split-3-GAL4: VT2064.p65ADZp@attP40; VT8168.ZpGAL4DBD@attP2

Split-4-GAL4: VT25965.p65ADZp@attP40; VT2064.ZpGAL4DBD@attP2

### Statistics

Mean values from behavioral experiments were compared using Krusakal-Wallis ANOVA or Mann-Whitney test. For Fisher’s exact test, two-tail p values were compared with controls. Statistical analyses were performed with GraphPad Prism software (SPSS Inc.).

## Supplemental Information

**Movie S1. Male-like social behaviors induced by activated *VT2064.GAL4 UAS-trpA1* females (Related to Figure 1)**

Videos of activated *VT2064.GAL4 UAS-trpA1* flies, showing specifically females displaying male-like courtship, aggression and chaining behaviors at ~28°C.

**Movie S2. *VT2064.GAL4* female induced male-like courtship without courtship song (Related to Figure 1)**

An audiovisual recording of an activated *VT2064.GAL4 UAS-trpA1* female displaying unilateral wing extension toward a wingless male without producing a courtship song at ~30°C.

**Movie S3. Male-like social behaviors induced by activated *Split-1-GAL4 UAS-trpA1* females (Related to Figure 2)**

Videos of activated *Split-1-GAL4 UAS-trpA1* females displaying male-like courtship, aggression and chaining behaviors at ~28°C.

**Movie S4. Position- and direction-selective following behavior in MARCM flies vs. non-selective following in *Split-1-GAL4 UAS-trpA1* (Related to Figure 3)**

Tracked videos, trajectories, angles and distances of activated MARCM Fly 1 (labeling one pC1-alpha neuron, blue arrow) showing following behavior specifically when a target fly (red arrow) moved from right to left across its visual field, and MARCM Fly 2 and a *Split-1-GAL4 UAS-trpA1* female showing position- and direction-selective following vs. non-selective following. Triangle and block represent start and end position of test (red) and target (blue) flies.

**Figure S1.**
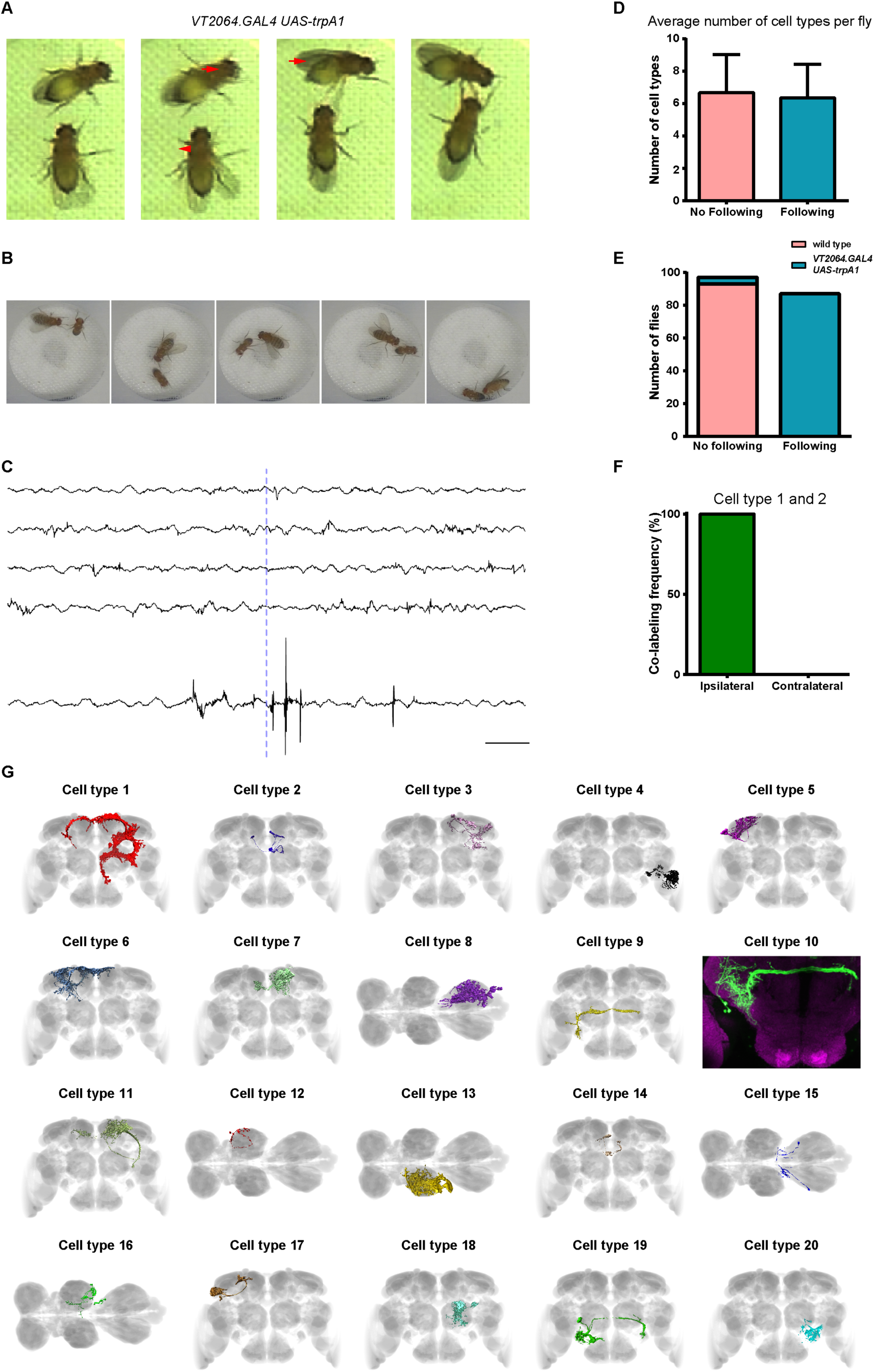
*VT2064.GAL UAS-trpA1* females induced silent courtship-like behavior and were analyzed using MARCM (Related to Figure 1) (A) A female carrying *VT2064.GAL4 UAS-trpA1*, upon activation, approached and extended one side of her wings (arrowhead), triggering rejection response from another female (arrows). (B and C) Images of unilateral wing extension displayed by *VT2064.GAL4 UAS-trpA1* females (B, left to right), and corresponding audio recordings (C, top to bottom). Wingless males were used as targets. Dashed red line indicates corresponding time points. Scale bar, 50ms. (D) Average of cell type numbers labeled in each MARCM fly in no following (n=34) and flowing groups (n=37). Data are represented as mean ± SD, with p=0.4522, Mann-Whitney test. (E) False negative (4.12%) and positive rates (0%) due to annotation of male-like following behavior were estimated by a blind control experiment using Canton S and *VT2064.GAL4 UAS-trpA1* flies, following a protocol with coded genotypes, predetermined test layouts, and delayed scoring (weeks later when memory faded). (F) Ipsilateral (n=37) and contralateral (n=0) colabeling frequency of cell type 1 and 2 in MARCM. (G) The 20 most frequently labeled cell types in MARCM.

**Figure S2.**
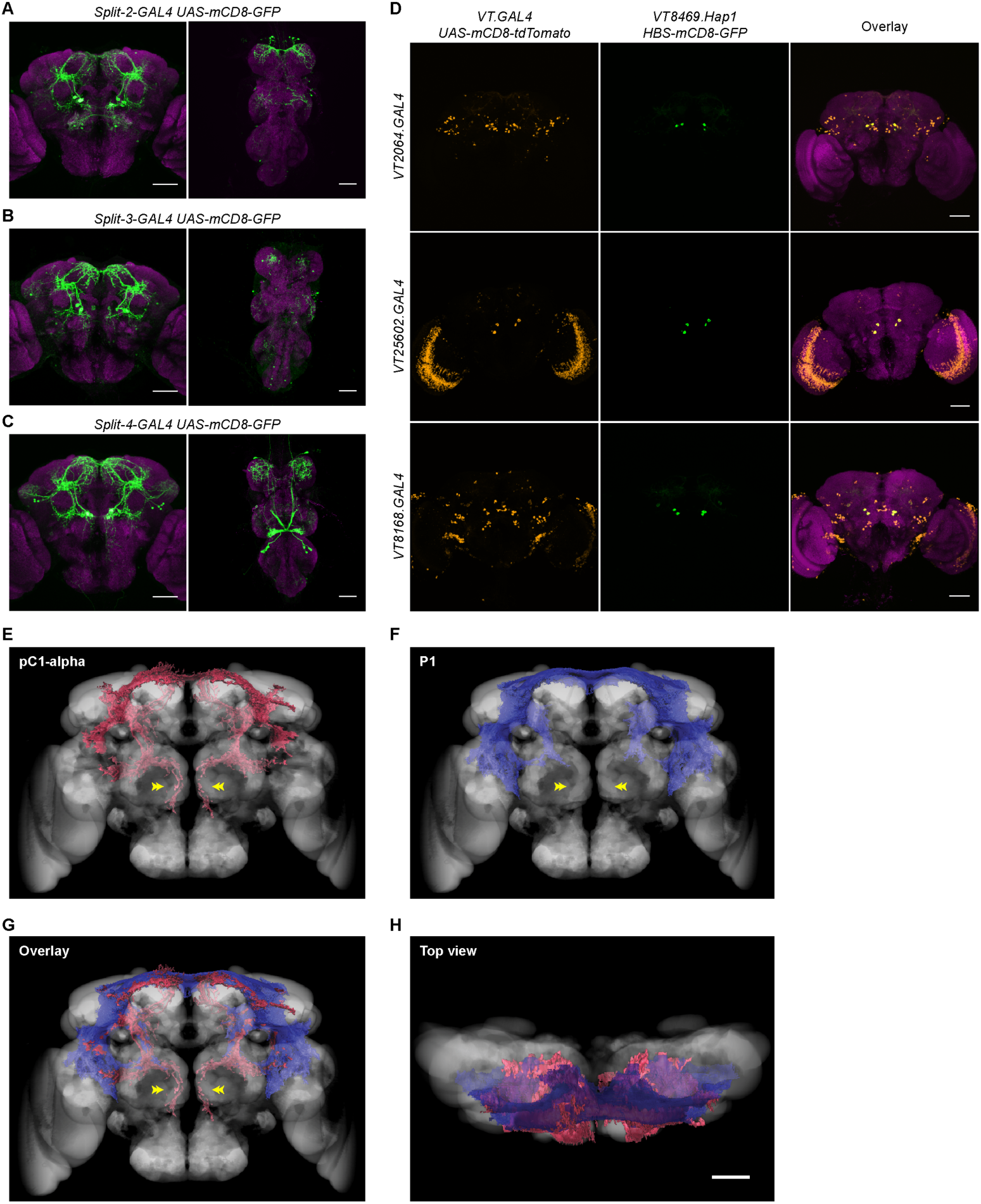
pC1-alpha is labeled by various driver lines and overlaps significantly with P1 (Related to Figure 2) (A to C) Brain (left) and VNC (right) of females carrying indicated split-GAL4 and *UAS-mCD8-GFP* transgenes, stained with anti-GFP (green) and nc82 (magenta). (D) Brains of flies carrying *VT8469.Hap1 HBS-mCD8-GFP* in combination with indicated GAL4 lines and *UAS-mCD8-tdTomato*, stained with anti-DsRed (orange), anti-GFP (green) and nc82 (magenta). (E-H) Registered segmentation of pC1-alpha neurons (E), P1 neurons (F), and their overlay (front view, G; top view, H) onto a common reference brain template. Double arrowheads indicate the characteristic medial ventral neurites of pC1-alpha.

